# *Prevotella copri*, a potential indicator for high feed efficiency in western steers

**DOI:** 10.1101/436832

**Authors:** Charles G. Brooke, Negeen Najafi, Katherine C. Dykier, Matthias Hess

**Affiliations:** Department of Animal Science, University of California, Davis, One Shields Ave, Davis CA, USA; California Department of Food and Agriculture, 1220 N Street, Sacramento, CA, USA

**Author notes:** Corresponding Author: Matthias Hess, 2251 Meyer Hall, Department of Animal Science, University of California, Davis, 1 Shields Ave, Davis, CA 95616, USA, P (530).

**Keywords:** animal microbiome, cattle, fecal microbiome, feed efficiency, microbial ecology, molecular marker

## Abstract

There has been a great interest to identify a microbial marker that can be used to predict feed efficiency of beef cattle. Such a marker, specifically one that would allow an early identification of animals with high feed efficiency for future breeding efforts, would facilitate increasing the profitability of cattle operations and simultaneously render them more sustainable by reducing their methane footprint. The work presented here suggests that *Prevotella copri* might be an ideal microbial marker for identifying beef cattle with high feed efficiency early in their life span and in the production cycle. Developing more refined quantification techniques that allow correlation of *P. copri* to feed efficiency of beef cattle that can be applied by lay people in the field holds great promise to improve the economy of cattle operations while simultaneously reducing their environmental impact by mitigating methane production from enteric fermentation.

## INTRODUCTION

Beef cattle is a significant source of protein in industrialized countries and feed constitutes up to 70% of the cost associated with meat production (1). In order to optimize these costs, there has been a great effort to identify and breed animals that possess higher feed efficiency. A common and widely accepted method of quantifying feed efficiency is to determine residual feed intake (RFI), which is a measure of the difference between an animal’s expected and actual feed intake (2) and is negatively correlated with the feed efficiency of an animal. Whereas methods for determining the RFI are well established, the biological and biochemical parameters that determine an animal’s RFI and therefore its feed efficiency phenotype are still poorly understood.

The rumen is host to a complex consortium of archaea, bacteria, fungi, and protozoa that work synergistically to degrade plant material and generate metabolites that deliver up to 80% of the energy of the host animal (3). Host genetics and diet impact the rumen microbiome of ruminant animals (4, 5) and even changes in the assemblage of the fiber-adherent population in response to the accumulation of metabolic intermediates has been reported (6). It has also been proposed that feed efficiency is linked to the host genetics and the microbiome assemblage within the gastrointestinal system of the animals (4, 7). Findings from studies investigating the correlation between microbiome members vary, which is not surprising since the experimental setup and analyses usually differ on multiple levels. A recent study focusing on the feed efficiency of dairy cows reported an inverse relationship between bacterial richness and feed efficiency and an increased abundance of *Megasphaera* sp in the highly efficient animals (8). A positive correlation (*p* < 0.02) between *Eubacterium* sp. and low feed efficiency was reported by Hernandez-Sanabria *et al.* for Hereford ×Aberdeen Angus steers that were fed a high energy diet (9), and Jami *et al.* reported a positive correlation between RFI and the uncultured rumen bacterium *RF39* [R = 0.51, p = 0.055,(10)]. Taxonomic markers at the family level associated with low efficient steers were identified recently (11), but the authors were not able to identify a taxonomic marker at higher resolution which could be used to predict the feed efficiency of the animals.

Since a microbial marker with sufficient resolution to enable a quick classification of animals into either feed efficiency category would be of great value to the cattle industry and have a direct positive impact on the environment by reducing the negative impact of cattle operations on methane emission, as well as reducing other waste products that are associated with maintaining large numbers of cattle, we investigated if such a marker would exist within samples found further along the gastrointestinal tract. Although collecting rumen fluid from fistulated cows or via tubulation is performed on a regular basis for scientific purposes, it is either not possible or practical on a cattle operation for the purpose of assigning feed efficiency to every animal that is considered for breeding. Since microbial community dynamics in response to changing diets in the reticulum have been reported to accurately reflect shifts within the rumen microbiome (12), we assessed if the fecal microbiome might contain microbial marker taxa that could be used for assigning feed efficiency to steers without investing significant amounts of resources. To identify potential microbial markers associated with feed efficiency phenotypes, we explored the fecal microbiome of low and high efficiency steers (based on RFI scores) by targeting the V4 region of the 16S rRNA gene.

## METHODS

### Feed Trial

A feeding trial involving 98 beef steers born fall 2014 was performed at UC Davis, CA. Animal procedures were performed under an animal care protocol IACUC # 17888. Residual feed efficiency was determined for each animal during a period of 56 days (07/06-08/31/15). Animals were fed *ad libitum* a diet composed of 69.8% flaked corn grain, 7.9% corn distillers dried grains, 5.9% wheat hay, 5.9% sudan hay, 5.6% liquid molasses, 1.17% urea, 1.8% yellow grease, 0.13% magnesium oxide, 1.42% calcium carbonate, 0.32% trace mineral salt, and 0.015% Rumensin^®^ (Elanco, Greenfield, IN). On August 31st, 36 animals were identified as either high or low RFI performers and will henceforth be referred to as low and high-efficiency steers respectively. Feed trial-meta data for sampled animals is provided in Supplementary Table S1.

### Sampling

Fecal samples were collected from 3 high efficiency and 3 low efficiency steers. Approximately 50 grams of fecal material was collected manually from 10-20 cm in the rectum and immediately flash frozen in liquid nitrogen. Samples were transported to the laboratory and stored at −20°C until further analysis.

### DNA extraction

DNA extraction was performed using the FastDNA SPIN Kit for Soil (MP Biomedicals, Solon, OH) according to the manufacturer’s protocol.

### PCR amplification, library preparation, and sequencing

The V4 region of the 16S rRNA gene was amplified using the 515F/806R primer set (13). For sequencing, forward and reverse sequencing oligonucleotides were designed. A detailed description of these oligonucleotide constructs and the conditions of the PCR reaction for amplification of the target region is provided in the Technical Appendix. Amplicon quantity was determined using a Qubit instrument with the Qubit High Sensitivity DNA kit (Invitrogen, Carlsbad, CA). Individual amplicon libraries were pooled, cleaned with Ampure XP beads (Beckman Coulter, Brea, CA), and sequenced using a 250 bp paired-end method (14) on an Illumina MiSeq at the DNA Technologies Core in the UC Davis Genome Center. Raw sequence reads were submitted to NCBI’s SRA under project accession number: PRJNA420377.

### Sequence Analysis

Forward and reverse sequencing oligonucleotides were designed to contain a unique 8 nt barcode (N), a primer pad (<u>underlined</u>), and the Illumina adaptor sequences (**bold**). Each sample was barcoded with a unique forward and reverse barcode combination. (Forward primer construct: **AATGATACGGCGACCACCGAGATCTACAC**-NNNNNNNN-<u>TATGGTAATT</u>-TGCCAGCMGCCGCGGTAA; reverse primer construct: **CAAGCAGAAGACGGCATACGAGAT**-NNNNNNNN-<u>AGTCAGTCAG</u>-GGACTACHVGGGTWTCTAAT.) Barcode combinations for each sample are provided in Supplementary Table S2. Each PCR reaction contained 1 Unit Kapa2G Robust Hot Start Polymerase (Kapa Biosystems, Boston, MA), 1.5 mM MgCl_2_, 10 pmol of each primer, and 1μL of DNA. The PCR was performed using the following conditions: 95°C for 2 min, followed by 30 cycles of 95°C for 10 s, 55°C for 15 s, 72°C for 15 s and a final extension of 72°C for 3 min. Sequencing resulted in a total of 298,445 reads, which were analyzed using Mothur v1.39.5 following the MiSeq SOP accessed on 09/01/2017. Using the *make.contigs* command, raw sequences were combined into contiguous sequences, which were filtered using *screen.seqs* to remove sequences that were >270 bp or contained ambiguous base calls to reduce PCR and sequencing error. Duplicate sequences were merged with *unique.seqs*, and the resulting unique sequences were aligned to the V4 region of the SILVA SEED alignment reference v123 (15) using *align.seqs*. Sequences were removed if they contained homopolymers longer than 8 bp or did not align to the correct region in the SILVA SEED alignment reference using *screen.seqs*. To further de-noise the data, sequences were pre-clustered within each sample allowing a maximum of 3 base pair differences between sequences using the *pre.cluster* command. Finally, chimeric sequences were identified using VSEARCH (16) and removed.

Quality filtered sequences were grouped into operational taxonomic units (OTUs) based on 97% sequence identity and classified using the Bayesian classifier and the Greengenes database (August 2013 release of gg_13_8_99) (17) with *classify.seqs*. Sequences that classified as mitochondria, chloroplasts, eukaryotes, or of unknown origin were removed using *remove.lineage.* Samples were rarefied to 7,012 sequences per sample, the smallest number of sequences across all collected samples. After pooling samples by feed efficiency, singleton and doubleton abundances were calculated with *filter.shared.* Chao1 diversity indices (18), Good’s coverage (19), Shannon indices (20), and inverse Simpson indices were calculated using *summary.single* to quantify coverage and alpha diversity for individual and pooled samples. Analysis of molecular variance (AMOVA) (21) was used to identify significant differences in community structure between feed efficiency phenotypes using the θ_YC_ distance matrix for the *amova* command (22), while linear discriminant analysis (LDA) effect size (LEfSe) (23) was used to identify indicator taxa that were significantly enriched in their respective groups.

### Statistical analysis

A two-tailed t-test was used to determine differences in diversity indices between high and low efficiency groups via SigmaPlot (SigmaPlot 11.0, Systat Software, Inc. San Jose, CA). Significant differences were defined as *p* < 0.05.

## RESULTS

### Sequencing and Quality Filtering

A total of 298,445 reads were generated from the fecal samples of 6 steers, with a mean of 21,170 reads per sample. After quality filtering and pooling the remaining 127,061 high quality reads into the two efficiency (high and low) groups, OTU based analysis (at 97% sequence identity) revealed 2,201 unique OTUs across the 6 samples. Of these 2201 OTUs, 1,510 (68.6%) were found in Low-efficiency steers and 1,471 (66.8%) were found in High-efficiency steers. Low and High-efficiency steers contained 730 and 691 exclusive OTUs respectively. Low and High-efficiency steers contained 469 and 534 singletons, and 224 and 234 doubletons, respectively. Singletons contributed to 0.56% (Low-efficiency) and 1.06% (High-efficiency) and doubletons contributed 0.27% and 0.53% of the quality-filtered reads respectively (Table 1).

**Table 1:** Statistics of 16S rRNA gene data. Data of steers were pooled based on their feed efficiency with n = 3 for each efficiency group.

|  | Low-efficiency | High-efficiency | Total |
| --- | --- | --- | --- |
| Raw Reads | 191,530 (100%) | 106,915 (100%) | 298,445 (100%) |
| Quality Filtered Reads | 82,851 (43%) | 44,172 (41%) | 127,061 (42%) |
| Doubletons | 224 | 234 |  |
| Singletons | 469 | 534 |  |
| Observed OTUs | 1,510 | 1,471 | 2,201 |
| Unique OTUs | 730 | 691 |  |
| Shared OTUs |  |  | 780 |
| Good's Coverage | 97% | 97% |  |

### Alpha and Beta Diversity Analysis

To estimate the microbial diversity within each group, rarefaction analyses were performed (Supplementary Fig. S1) and species richness as well as diversity indices were calculated (Table 2). No difference in Chao1, inverse Simpson or Shannon indices were detected (p > 0.05) between low and high efficiency groups. Good’s coverage estimates were ≥97% in all samples (Table 2), suggesting that sequencing efforts were sufficient to recover a large proportion of the microbial diversity in each of the samples under investigation.

**Table 2:**
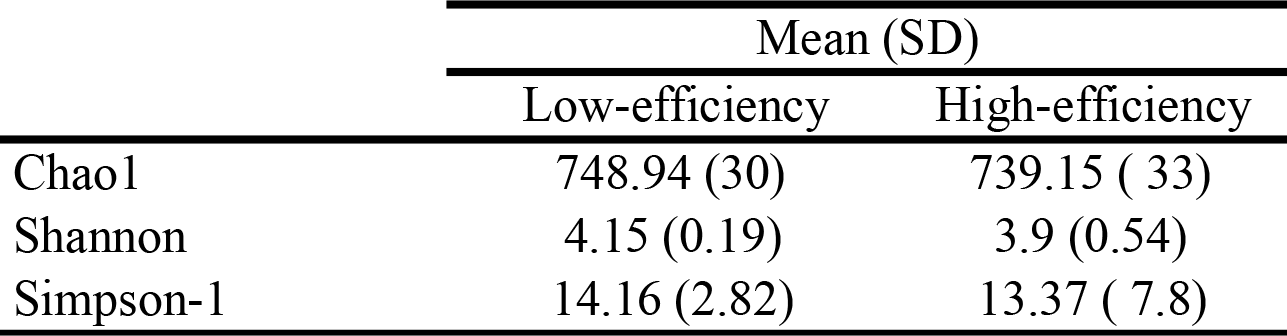
Diversity indices of 16S rRNA gene sequences from animals of low and high efficiency groups (n = 3 for each group)

Variance of the microbial community between and among low and high efficiency steers were quantified using a θ_YC_ distance matrix (22) and subsequent AMOVA and HOMOVA analyses. The microbial community did not differ significantly between the two groups of steers (AMOVA, *p* >*0.05*), and the variance between the two groups was not significantly different (HOMOVA, *p* >*0.05*).

### Microbial Community Structure

Across all samples, 2,201 OTUs were classified into one archaeal and 21 bacterial phyla. The 10 most abundant phyla recruited >98% of the reads generated from the microbial communities of both low and high efficiency groups (Fig. 1). *Firmicutes* dominated the microbial communities of both phenotypes and recruited on average 67.3% (± 6%) and 51% (±27%) of the reads in the low and high efficiency groups respectively. *Bacteroidetes* was the second most dominant phylum in both groups with a similar average percentage of 20% (±5%) and 19.3% (±3%) of the total reads. The remaining sequences from the low and high efficiency groups were assigned to the *Proteobacteria* [0.4% (±0.08%) and 5% (±7%)], *Tenericutes* [0.5%(±0.32%) and 0.3%(±0.18%)], *Euryarcheota* [3%(±4%) and 3%(±4%)], *Actinobacteria* [0.5% (±0.42%) and 0.6% (±0.37%)], *Spirochetes* [4.6% (±6%) and 2.4%(±3%)] and *Verrucomicrobia* [0.7%(±1%) and 0.8%(±0.7%)]. *Fusobacteria* were largely associated with the high efficiency phenotype, with an average of 16% of the total reads, with an increased proportion (46%) of reads in one subject (animal #61) of the high efficiency phenotype. This portion of reads was attributed to a single OTU that was classified as *Cetobacteria somerae*. The average relative abundance of *Proteobacteria* was higher in the high efficiency group recruiting 0.4% (±0.08%) and 5% (±7%) in low and high efficiency animals respectively. The amplicon library of animal #61 drove this difference as well, with *Proteobacteria* representing 13% of the total reads from this sample. The differential abundance of proteobacteria in this sample can also be attributed to a single OTU, classified to the *Succinovibrionaceae* family.

**Figure 1:**
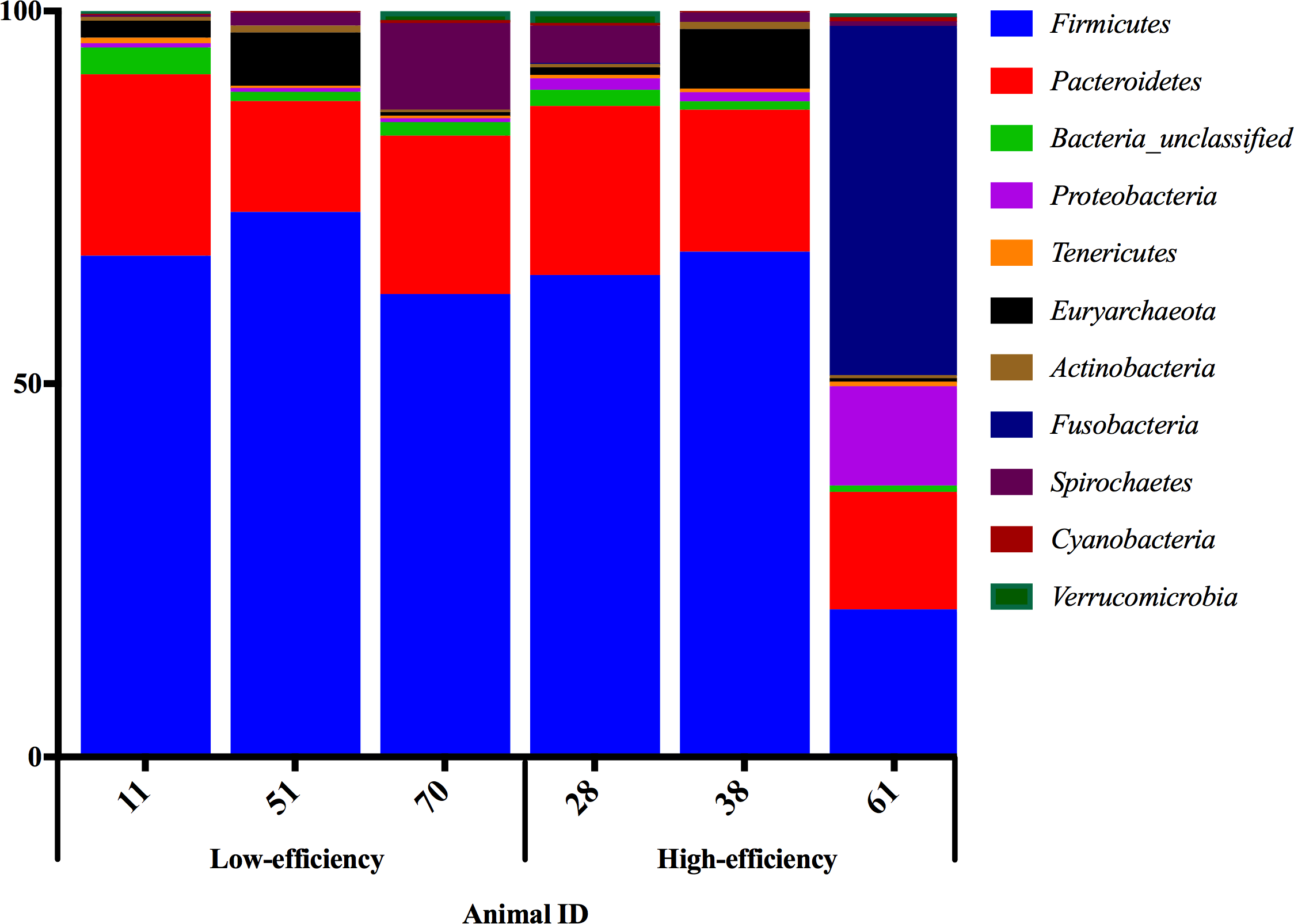
Phylum level composition of fecal microbiome.

### Indicator taxa

Overall, 24 taxa at various phylogenetic levels were differentially abundant in their respective samples (LDA scores >2.0, *p* value ≤0.05, Table 3). To facilitate data interpretation and subsequent discussion we focused on the most highly enriched taxa, reporting only taxa with LDA scores >3. Four OTUs with LDA scores >3 were identified as indicators of the low efficiency group and three were identified in the high efficiency group. An OTU in the *Ruminococcaceae* family generated the highest LDA score (4.67) in the low efficiency group with a mean read percentage of 23.5% (± 1.6%) compared to the high efficiency group with 14% (± 5.4%) of the total reads. An OTU that mapped to the genus *Clostridium* of the *Firmicutes* phylum was assigned the second highest LDA score within the low efficiency group (3.63) accounting for an average of 1.32% (±0.6%) of the reads for the low efficiency group and 0.5% (±0.05%) of the high efficiency reads. The remaining two OTUs that generated LDA scores >3 and classified as indicators of the low efficiency group were: An OTU that classified to the *Oscillispira* genera [LDA score = 3.31, low efficiency group abundance = 0.48% (±0.12%), High-efficiency = 0.27% (±0.14%)] and an OTU that was classified to the *Bacteriodaceae* family [LDA score = 3.01, low efficiency abundance = 0.44% (±0.23%), low efficiency group abundance = 0.31% (±0.21%)]. Three OTUs with LDA scores >3 were more abundant in the high efficiency group. *Prevotella copri* was assigned an LDA score of 4.14 and on average, accounted for 3% (±0.6%) of the reads in the high efficiency group, and 0.14% (±0.02%) in the low efficiency group. An OTU classified as member of the genus *Prevotella* had the second highest LDA score within the high efficiency group [LDA score = 3.32, high efficiency group abundance = 0.53% (±0.2%), low efficiency abundance = 0.35% (±0.2%)]. The Last OTU in the high efficiency group with an LDA score >3 classified to the *Paraprevotella* genus [LDA score= 3.28, high efficiency = 0.09% (±0.09%), low efficiency = 0.07% (±0.07%)]

**Table 3.** Operational Taxonomic Units (OTUs) identified as indicator taxa for low and high efficiency group based on linear discriminant analysis.

| OTU | Taxonomic Classification | LDA Score | % of Total Reads ( ±STD) | | Efficiency Group |
| --- | --- | --- | --- | --- | --- |
|  |  |  | Low-efficiency | High-efficiency |  |
| Otu0001 | f__Ruminococcaceae_unclassified(100) | 4.67 | 23.56 (1.6) | 14.22 (5.4) | Low |
| Otu0015 | g__Prevotella; s__copri(100) | 4.14 | 0.14 (0.02) | 3.01 (0.6) | High |
| Otu0016 | g__Clostridium_unclassified(60) | 3.63 | 1.33 (0.63) | 0.46 (0.06) | Low |
| Otu0028 | g__Prevotella_unclassified(100) | 3.32 | 0.35 (0.20) | 0.53 (0.2) | Low |
| Otu0034 | g__Oscillospira_unclassified(100) | 3.31 | 0.48 (0.12) | 0.27 (0.14) | Low |
| Otu0114 | g__Paraprevotella_unclassified(100) | 3.28 | 0.07 (0.07) | 0.09 (0.09) | High |
| Otu0032 | f__Bacteroidaceae_unclassified(100) | 3.01 | 0.44 (0.23) | 0.31 (0.21) | Low |
| Otu0128 | f__Lachnospiraceae_unclassified(77) | 2.98 | 0.06 (0.02) | 0.10 (0.14) | High |
| Otu0087 | g__Ruminococcus; s__bromii(99) | 2.93 | 0.18 (0.01) | 0.15 (0.07) | Low |
| Otu0109 | g__5-7N15_unclassified(100) | 2.78 | 0.14 (0.06) | 0.10 (0.06) | Low |
| Otu0170 | g__Prevotella_unclassified(100) | 2.78 | 0.09 (0.05) | 0.02 (0.02) | Low |
| Otu0191 | g__Dorea_unclassified(99); | 2.71 | 0.05 (0.03) | 0.05 (0.03) | Low |
| Otu0180 | g__Ruminococcus_unclassified(100) | 2.67 | 0.10 (0.05) | 0.02 (0.02) | Low |
| Otu0240 | f__Lachnospiraceae_unclassified(100) | 2.64 | 0.03 (0.02) | 0.06 (0.05) | High |
| Otu0164 | f__Lachnospiraceae_unclassified(100) | 2.59 | 0.10 (0.04) | 0.05 (0.03) | Low |
| Otu0236 | C__L635J-21_unclassified(100) | 2.53 | 0.01 (0.01) | 0.13 (0.11) | High |
| Otu0264 | g__Oscillospira_unclassified(100) | 2.51 | 0.02 (0.004) | 0.05 (0.03) | Low |
| Otu0417 | f__Ruminococcaceae_unclassified(100) | 2.40 | 0 (0) | 0.02 (0.02) | Low |
| Otu0280 | f__Lachnospiraceae_unclassified(98) | 2.34 | 0.08 (0.04) | 0.005 (0.004) | Low |
| Otu0267 | g__Anaerovorax_unclassified(100) | 2.34 | 0.06 (0.03) | 0.01 (0.005) | High |
| Otu0310 | f__Lachnospiraceae_unclassified(85) | 2.31 | 0.03 (0.02) | 0.01 (0.01) | Low |
| Otu0552 | g__Prevotella_unclassified(100) | 2.18 | 0 (0) | 0.05 (0.05) | Low |
| Otu0487 | g__Anaeroplasma_unclassified(100) | 2.16 | 0.005 (0.005) | 0 (0) | Low |
| Otu0382 | p__Bacteroidetes_unclassified(100) | 2.08 | 0.02 (0.01) | 0.01 (0.01) | Low |

## DISCUSSION

Results obtained from 16S rRNA-based amplicon sequencing suggests that the fecal microbiomes of steers with low and high feed efficiency are not significantly different at the community level (p > 0.05). This coincides with the previous observations by Myer et al (2015) who investigated the fecal microbiome of 32 steers classified by a Cartesian coordinate system of average daily gain and average daily feed intake (24). Taking our findings and results from previous studies into consideration, it appears 16S rRNA amplicon sequencing does not lend itself as a rapid screening tool to classify individual animals regarding their feed efficiency at the phylum level. However, when comparing rumen samples of steers with high and low feed efficiency at the family and genus level, differences in their microbiome assemblage can be detected (24, 25). Although these differences might be of great value to the scientific community for efficiency related studies, acquiring rumen samples from a large number of animals and performing time consuming 16S rRNA sequencing appears unlikely to be an option for the average cattle producer who needs reliable information regarding the feed efficiency of an individual animal within a few days if not even within hours. Microorganisms classified at the species level could provide such a screenable marker, if they are enriched significantly enough in one of the efficiency groups and found in sufficient abundance in the feces of a steer to warrant detection via basic but still reliable molecular techniques such as qPCR. Our data indicate that *Prevotella copri* was enriched in the fecal microbiome of high-efficiency steers. This data suggests that *P. copri* might be a potential marker for increased feed efficiency in the feces of beef cattle. It should be noted that while Shabat et al (2016) did not specifically associate *P.copri* with high efficiency dairy cattle, sequencing data from high efficiency animals contained reads derived from *P. copri* (11). The finding that *Prevotella copri* is associated with animals of high feed efficiency is not surprising, since *P. copri* is capable of utilizing a wide variety of carbohydrates, and has been identified before as potential key factor in shaping gut function and host health. In a study encompassing both humans and mice, *P.copri* was shown to enhance the ability to utilize complex polysaccharides (26), increase glycogen storage capability, augment glucose tolerance in mice and to be associated with insulin resistance (27, 28), a major factor in weight gain. Also Myer at al (24) detected a significant enrichment of *P. copri* in fecal samples of steers with high daily gain and intake, but the relative abundance of *P. copri* within the microbiome from these steers was reflecting the relative abundance (<0.01%) we observed in steers with low feed efficiency. This might be attributed to the choice of primer set that was used to generate the data or to the fact that steers deemed high efficiency would have been classified as low efficiency animals in our study. Here we utilized primers targeting the V4 region, which have shown to enable microbiome profiles at an increased resolution (29) and improved reproducibility (30). Our analysis revealed a 20-fold increase in the abundance of *P. copri*, with 0.14% and 3% of the reads from the low and high feed efficient microbiomes respectively. Recently, it was shown that the methods for DNA preparation have a significant affect on microbiome data (31). Specifically, the kit utilized for the study presented here had a positive association with the Prevotellaceae family, which may have aided in the identification of *P. copri*.

In addition to increasing the profitability of a cattle operation from the standpoint of increased weight gain per unit of feed, increased feed efficiency also decreases the environmental footprint of a cattle operation by reducing methane production and release from a feed efficient animal due to reduced dry matter intake (32). The exact molecular mechanisms of reduced methane emission in more efficient ruminants are still not fully understood, but it is possible that hydrogen redistribution plays a significant role, since methane production can be depleted in the presence of a competing hydrogen sink such as the volatile fatty acid propionate (33). Some *Bacteriodetes*, like *P. copri*, produce succinate, the precursor to propionate, as their predominate fermentation byproduct, which in addition to its potential impact on gluconeogenesis (27), may inhibit methane synthesis. Whether animal genetics impact the relative abundance of this organism is yet to be determined. Future work investigating the abundance dynamics of *P.copri* throughout the life of animals with different genetic backgrounds may provide valuable data to fill this knowledge gap. It would be beneficial to develop approaches that would allow the classification of animals with low or high feed efficiency in an early life stage so that informed decisions could be made regarding breeding practices. This study characterizes *Prevotella copri* as a potential indicator of feed efficiency from the fecal microbiome of western steers and should stimulate directed research into the dynamics of this organism through the life of beef cattle as well as the fecal microbiome as a whole.

## ACKNOWLEDGEMENTS

We thank Dr. Roberto Sainz for the fecal samples and his comments on the manuscript. We also thank Yuki Okatsu for helping with sample collection and DNA extraction.

## REFERENCES

1. Kahn L, Cottle D. Beef cattle production and trade: Csiro Publishing; 2014.

2. Koch RM, Swiger LA, Chambers D, Gregory KE. Efficiency of Feed Use in Beef Cattle1. Journal of Animal Science. 1963;22(2):486–94.

3. Wolin MJ. The Rumen Fermentation: A Model for Microbial Interactions in Anaerobic Ecosystems. In: Alexander M, editor. Advances in Microbial Ecology: Volume 3. Boston, MA: Springer US; 1979. p. 49–77.

4. Myer PR, Wells JE, Smith TPL, Kuehn LA, Freetly HC. Gut bacterial communities and their association with production parameters in beef cattle. J Anim Sci. 2016;94(suppl_2):183–4.

5. Henderson G, Cox F, Ganesh S, Jonker A, Young W, Global Rumen Census C, et al. Rumen microbial community composition varies with diet and host, but a core microbiome is found across a wide geographical range. Sci Rep. 2015;5:14567.

6. Piao H, Lachman M, Malfatti S, Sczyrba A, Knierim B, Auer M, et al. Temporal dynamics of fibrolytic and methanogenic rumen microorganisms during in situ incubation of switchgrass determined by 16S rRNA gene profiling. Frontiers in Microbiology. 2014;5:307.

7. Roehe R, Dewhurst RJ, Duthie CA, Rooke JA, McKain N, Ross DW, et al. Bovine Host Genetic Variation Influences Rumen Microbial Methane Production with Best Selection Criterion for Low Methane Emitting and Efficiently Feed Converting Hosts Based on Metagenomic Gene Abundance. PLoS Genet. 2016;12(2):e1005846.

8. Shabat SK, Sasson G, Doron-Faigenboim A, Durman T, Yaacoby S, Berg Miller ME, et al. Specific microbiome-dependent mechanisms underlie the energy harvest efficiency of ruminants. Isme J. 2016;10(12):2958–72.

9. Hernandez-Sanabria E, Goonewardene LA, Wang Z, Durunna ON, Moore SS, Guan LL. Impact of Feed Efficiency and Diet on Adaptive Variations in the Bacterial Community in the Rumen Fluid of Cattle. Applied and Environmental Microbiology. 2012;78(4):1203–14.

10. Jami E, White BA, Mizrahi I. Potential Role of the Bovine Rumen Microbiome in Modulating Milk Composition and Feed Efficiency. PLOS ONE. 2014;9(1):e85423.

11. Li F, Zhou M, Ominski K, Guan LL. Does the rumen microbiome play a role in feed efficiency of beef cattle?1. J Anim Sci. 2016;94(suppl_6):44–8.

12. Kim M, Kim J, Kuehn LA, Bono JL, Berry ED, Kalchayanand N, et al. Investigation of bacterial diversity in the feces of cattle fed different diets1. Journal of Animal Science. 2014;92(2):683–94.

13. Caporaso JG, Lauber CL, Walters WA, Berg-Lyons D, Lozupone CA, Turnbaugh PJ, et al. Global patterns of 16S rRNA diversity at a depth of millions of sequences per sample. Proceedings of the National Academy of Sciences. 2011;108(Supplement 1):4516–22.

14. Kozich JJ, Westcott SL, Baxter NT, Highlander SK, Schloss PD. Development of a Dual-Index Sequencing Strategy and Curation Pipeline for Analyzing Amplicon Sequence Data on the MiSeq Illumina Sequencing Platform. Applied and Environmental Microbiology. 2013;79(17):5112–20.

15. Quast C, Pruesse E, Yilmaz P, Gerken J, Schweer T, Yarza P, et al. The SILVA ribosomal RNA gene database project: improved data processing and web-based tools. Nucleic Acids Research. 2013;41(Database issue):D590–D6.

16. Rognes T, Flouri T, Nichols B, Quince C, Mahé F. VSEARCH: a versatile open source tool for metagenomics. PeerJ. 2016;4:e2584.

17. DeSantis TZ, Hugenholtz P, Larsen N, Rojas M, Brodie EL, Keller K, et al. Greengenes, a Chimera-Checked 16S rRNA Gene Database and Workbench Compatible with ARB. Applied and Environmental Microbiology. 2006;72(7):5069–72.

18. Chao A. Nonparametric Estimation of the Number of Classes in a Population. Scandinavian Journal of Statistics. 1984;11(4):265–70.

19. Good IJ. The Population Frequencies of Species and the Estimation of Population Parameters. Biometrika. 1953;40(3/4):237–64.

20. Shannon C. À mathematical theory of communication. Bell System Technology. Journal. 1948.

21. Excoffier L, Smouse PE, Quattro JM. Analysis of molecular variance inferred from metric distances among DNA haplotypes: application to human mitochondrial DNA restriction data. Genetics. 1992;131(2):479–91.

22. Yue JC, Clayton MK. A similarity measure based on species proportions. Communications in Statistics-Theory and Methods. 2005;34(11):2123–31.

23. Segata N, Izard J, Waldron L, Gevers D, Miropolsky L, Garrett WS, et al. Metagenomic biomarker discovery and explanation. Genome biology. 2011;12(6):R60.

24. Myer PR, Smith TP, Wells JE, Kuehn LA, Freetly HC. Rumen microbiome from steers differing in feed efficiency. PLoS One. 2015;10(6):e0129174.

25. Li F, Guan LL. Metatranscriptomic profiling reveals linkages between the active rumen microbiome and feed efficiency in beef cattle. Appl Environ Microbiol. 2017.

26. Kovatcheva-Datchary P, Nilsson A, Akrami R, Lee YS, De Vadder F, Arora T, et al. Dietary Fiber-Induced Improvement in Glucose Metabolism Is Associated with Increased Abundance of Prevotella. Cell Metab. 2015;22(6):971–82.

27. De Vadder F, Kovatcheva-Datchary P, Zitoun C, Duchampt A, Bäckhed F, Mithieux G. Microbiota-Produced Succinate Improves Glucose Homeostasis via Intestinal Gluconeogenesis. Cell Metabolism. 2016;24(1):151–7.

28. Pedersen HK, Gudmundsdottir V, Nielsen HB, Hyotylainen T, Nielsen T, Jensen BAH, et al. Human gut microbes impact host serum metabolome and insulin sensitivity. Nature. 2016;535(7612):376–81.

29. Soergel DAW, Dey N, Knight R, Brenner SE. Selection of primers for optimal taxonomic classification of environmental 16S rRNA gene sequences. ISME J. 2012;6(7):1440–4.

30. Tremblay J, Singh K, Fern A, Kirton ES, He S, Woyke T, et al. Primer and platform effects on 16S rRNA tag sequencing. Frontiers in microbiology. 2015;6.

31. Vaidya JD, van den Bogert B, Edwards JE, Boekhorst J, van Gastelen S, Saccenti E, et al. The Effect of DNA Extraction Methods on Observed Microbial Communities from Fibrous and Liquid Rumen Fractions of Dairy Cows. Front Microbiol. 2018;9:92.

32. Basarab JA, Beauchemin KA, Baron VS, Ominski KH, Guan LL, Miller SP, et al. Reducing GHG emissions through genetic improvement for feed efficiency: effects on economically important traits and enteric methane production. Animal. 2013;7(Suppl 2):303–15.

33. Janssen PH. Influence of hydrogen on rumen methane formation and fermentation balances through microbial growth kinetics and fermentation thermodynamics. Animal Feed Science and Technology. 2010;160(1):1–22.

